# The CDK7 inhibitor CT7001 (Samuraciclib) targets proliferation pathways to inhibit advanced prostate cancer

**DOI:** 10.1101/2022.06.29.497030

**Authors:** Theodora A. Constantin, Anabel Varela-Carver, Kyle K. Greenland, Gilberto Serrano de Almeida, Lucy Penfold, Simon Ang, Alice Ormrod, Edward K. Ainscow, Ash K. Bahl, David Carling, Matthew J. Fuchter, Simak Ali, Charlotte L. Bevan

## Abstract

**Background:** Current strategies to inhibit the androgen receptor (AR) are circumvented in castration-resistant prostate cancer (CRPC). Cyclin-dependent kinase 7 (CDK7) promotes AR signalling, in addition to established roles in cell cycle and global transcription regulation, together, providing a rationale for its therapeutic targeting in CRPC.

**Methods:** The antitumour activity of CT7001, an orally bioavailable CDK7 inhibitor, was investigated across CRPC models *in vitro* and in xenograft models *in vivo*. Cell-based assays and transcriptomic analyses of treated xenografts were employed to investigate the mechanism driving activity of CT7001, alone and in combination with the antiandrogen enzalutamide.

**Results:** CT7001 selectively engages with CDK7 in prostate cancer cells, causing inhibition of proliferation and cell cycle arrest. Activation of p53, induction of apoptosis, and suppression of transcription mediated by full-length and constitutively active AR splice variants contribute to antitumour efficacy *in vitro*. Oral administration of CT7001 represses growth of CRPC xenografts and significantly augments growth inhibition achieved by enzalutamide. Transcriptome analyses of treated xenografts indicate cell cycle and AR inhibition as the mode of action of CT7001 *in vivo*.

**Conclusions:** This study supports CDK7 inhibition as a strategy to target deregulated cell proliferation and demonstrates CT7001 is a promising CRPC therapeutic, alone or in combination with AR-targeting compounds.

## BACKGROUND

Prostate cancer is the second most common malignancy in men worldwide and a leading cause of cancer-related death [1]. The androgen receptor (AR) is a ligand-dependent nuclear receptor that plays a critical role in prostate cancer initiation and progression [2]. Therapies aiming to ablate AR activity by preventing ligand-dependent receptor activation are initially effective, but circumvented following progression to a therapy-resistant stage termed castration-resistant prostate cancer (CRPC). Commonly, castration resistance mechanisms facilitate reactivation of the AR signalling pathway and downstream transcriptional programs [3–5]. One phenomenon associated with disease progression, the expression of AR splice variants (AR-V), has drawn interest as a potential mechanism mediating androgen independency in CRPC [6–8]. AR-Vs lack the AR ligand-binding domain (LBD) but retain the N-terminal activation function, thereby having the potential to mediate AR signalling while being insensitive to approved AR-targeted therapies, all of which target the LBD [9]. Novel therapeutic approaches are therefore needed for the treatment of CRPC, with strategies that interfere with oncogenic AR transcription in a LBD-independent manner being of particular interest [10].

Cyclin-dependent kinase 7 (CDK7) is a recognised anticancer drug target due to its regulatory roles in cell division and transcription [11, 12]. As part of the CDK-activating kinase (CAK) complex together with cyclin H and MAT1, CDK7 phosphorylates the T-loop of cell cycle CDKs, including CDK1, -2, -4 and -6, stimulating their activities in a temporal manner to drive progression through the cell cycle [13–16]. CAK also functions as a component of the general transcription factor TFIIH, enabling transcription initiation by phosphorylating serine-5 residues (S5) in the RNA polymerase II (PolII) C-terminal domain (CTD) [17].

Although phosphorylation of PolII CTD is essential for mRNA synthesis [18], whether CDK7 activity is strictly required for basal transcription is still debated [19–22]. It has previously been shown that CDK7 phosphorylates AR at serine-515 (S515) to regulate transactivation and proteasomal degradation of AR, and facilitates assembly of a transcriptionally active AR-coregulator complex [23–25]. Thus, CDK7 may be promoting oncogenic AR transcription in CRPC by both AR-specific and more global transcriptional effects. In line with a tumour-promoting role in CRPC, high tumour CDK7 protein levels are associated with faster biochemical recurrence, marked by rises in serum prostate-specific antigen (PSA) [26].

The multiple roles of CDK7 commend it as a drug target for CRPC. However, a fundamental understanding of the mechanistic effects of selective CDK7 inhibition in CRPC is currently lacking. In a previous study, the covalent CDK7 inhibitor THZ1 was used [23], however, this also targets CDK12/13, which substantially obscures the contribution of CDK7 inhibition to its downstream effects [21, 22, 27]. CT7001/Samuraciclib (formerly ICEC0942) is a novel, orally bioavailable, ATP-competitive inhibitor of CDK7 with preclinical activity in models of breast cancer, colon cancer, and acute myeloid leukaemia [28–30]. CT7001 is currently in phase I/II clinical trials for advanced solid malignancies (NCT03363893). Recently released data from this study demonstrated acceptable safety profile and evidence of antitumour activity, including PSA reductions in 4 CRPC patients [31]. Notably, the combination of CT7001 with fulvestrant was well-tolerated and reduced tumour burden in a cohort of patients with CDK4/6 inhibitor-resistant/hormone receptor-positive/HER2-negative breast cancer [32]. As CT7001 is advancing rapidly through clinical evaluation, a more comprehensive investigation of its preclinical activity in prostate cancer is warranted. Here, we describe the mechanism of tumour inhibition achieved by CT7001 in prostate cancer models *in vitro* and *in vivo* and explore its efficacy as monotherapy and in combination with the widely used AR antagonist enzalutamide.

## MATERIALS & METHODS

### Chemicals

CT7001 was provided by Carrick Therapeutics. Apalutamide, enzalutamide, darolutamide and bavdegalutamide were purchased from MedChemExpress. Mibolerone was purchased from PerkinElmer.

### Cell lines

LNCaP, C4-2, VCaP, DU145, PC3, PNT1A, BPH-1 cell lines were obtained from the ATCC as frozen stocks. 22Rv1 and 22Rv1 FL-AR KO were a gift from Dr Luke Gaughan (Newcastle University) [33]. C4-2B cells were a gift from Prof Ian Mills (Oxford University) [34]. LNCaP/Luc cells are stably transfected with an AR-specific luciferase reporter construct [35] and selection was maintained with 500μg/mL G418 (Sigma) and 10μg/mL blasticidin (Melford Biolaboratories). All cell lines were cultured according to ATCC recommendations in growth medium with 10% foetal calf serum (FCS) and 2mM L-glutamine at 37°C in humidified incubators maintained at 5% CO_2_. Cell line authenticity was confirmed using short tandem repeat analysis performed through the MWG Eurofins Human Cell Line Authentication service. All cell lines were regularly tested for Mycoplasma infection using the MycoAlert Detection Kit (Lonza).

### Cellular thermal shift assay (CeTSA) in intact LNCaP cells

The melting temperature (T_m_) of CDK targets were determined using a Boltzmann sigmoidal curve (least squares fit) fitted through the average normalised data points from all human lines available in the Meltome atlas [36]. The melt curve for CDK7 was generated as previously described [37]. Isothermal dose-response fingerprints were generated at 54°C as previously described [37] from intact LNCaP cells treated with dimethylsulfoxide (DMSO) or CT7001 (concentrations range 0.078-20μM) for 3 hours..

### Protein extraction

Following incubation with fresh growth medium containing CT7001, cells were harvested and lysed as previously described [35]. Protein content was measured using the Pierce BCA Protein Assay Kit (ThermoFisher).

### Immunoblotting

Immunoblotting was carried out as previously described [38] using the following primary antibodies: CDK7 (Cell Signaling Technology, 2916), CDK1 (Cell Signaling Technology, 9116), CDK2 (Cell Signaling Technology, 2546), CDK4 (Cell Signaling Technology, 12790), CDK9 (Cell Signaling Technology, 2316), β-actin (Abcam, ab6276), GAPDH (Cell Signaling Technology, 2118), P-S5 PolII (Abcam, ab5401), P-S2 PolII (Abcam, ab5095), PolII (Abcam, ab817), P-S780 Rb (Abcam, ab47763), P-S807/811 Rb (Cell Signaling Technology, 8516), Rb (Abcam, ab6075), cyclin H (Abcam, ab54903), MAT1 (Santa Cruz Biotechnology, sc135981), P-S15 p53 (Cell Signaling Technology, 9284), p53 (Santa Cruz Biotechnology, sc-126), P-T161/T160 CDK1/CDK2 (Abcam, ab201008), P-S10 Histone H3 (Abcam, ab139417), AR (MilliporeSigma, 06-680), P-T1457 MED1 (Abcam, ab60950), MED1 (Bethyl Laboratories, A300-793A), cleaved PARP1 (Abcam, ab4830), p21 (Santa Cruz Biotechnology, sc-817).

### Cell growth and drug synergy assays

Cell number was assayed using the Sulforhodamine B (SRB) assay as previously described [39]. To determine growth rate (GR) values for each treatment and derive GR metrics, a file containing cell number data from day 0 and day 3 for was uploaded on the GRcalculator platform (www.grcalculator.org/grcalculator) [40].

CT7001 combinations with AR-targeted therapies were tested in a 6×5 matrix in LNCaP, C4-2B, and PC3 cells. Cell numbers on day 0 and 3 were assayed using the SRB assay. Percent growth inhibition was calculated relative to vehicle control using day 3 values corrected by subtracting day 0 values. A file containing growth inhibition (%) data for each cell line was uploaded on the SynergyFinder 2.0 web application (<u>synergyfinder.fimm.f</u>i) [41]. Bliss independence scores were calculated using default parameters.

### Caspase 3/7 assay

Caspase 3/7 activation assays were performed using the Caspase-Glo 3/7 Assay System (Promega), 72 hours after treatment with CT7001. Luminescence was measured using the VICTOR Light luminometer (PerkinElmer). Signal was normalised to cell number on day 3 measured by SRB assay.

### RT-qPCR and TaqMan Low Density Array

Total RNA was extracted from treated cells using the RNeasy Mini kit (QIAGEN). cDNA was synthesized from 1μg of RNA with oligo-dT primers using RevertAid First Strand cDNA Synthesis Kit (ThermoFisher). Quantitative Polymerase Chain Reaction (qPCR) was performed in a QuantStudio 7 Flex Real-Time PCR System (Applied Biosystems) using SYBR Green Real-Time PCR Master mix (Invitrogen) and primers designed using Primer-BLAST (listed in **Additional File 2** – **Supplementary Table 1**). Specificity was validated using melt curve analysis. Gene expression was normalised to RPL19, GAPDH and BACTIN and data were analysed using the ΔΔCt method.

TaqMan Low-Density Array microfluidic cards were used to assay the expression of 32 AR targets (a list of the TaqMan qPCR assays is provided in **Additional File 2, Supplementary Table 2**). 22Rv1 FL-AR KO cells were treated 1μM CT7001 in androgen-depleted medium (phenol red-free RPMI-1640 with 5% charcoal-stripped FCS). LNCaP and VCaP cells were treated with 1μM CT7001 in androgen-repleted medium (androgen-depleted medium supplemented with 10nM mibolerone). Total RNA was extracted after 24 hours, and cDNA was synthesized as above. cDNA was mixed with TaqMan Gene Expression Master Mix (ThermoFisher) and loaded on the microfluidic card. Gene expression was normalised to GAPDH, RPLP0, TBP, and 18S and the data were analysed using the ΔΔCt method.

### Cell cycle analysis

Asynchronous cells treated with DMSO or CT7001 for 72 hours were harvested and fixed in 70% ice-cold ethanol overnight. Fixed cells were washed twice with PBS, stained with the Muse Cell Cycle Reagent (Luminex), and analysed using the Guava Muse Cell Analyzer (Luminex).

### Plasmids and site-directed mutagenesis

The following plasmids have been previously described: pSV-AR coding human AR, TAT-GRE-E1B-LUC coding AR-responsive luciferase reporter, and BOS-β-galactosidase coding a constitutive reporter [42].

Plasmid constructs harbouring AR splice variants (pCMV5-CE1 coding AR-V1, pCMV5-CE2 coding AR-V6, pCMV5-CE3 coding AR-V7, pCMV5-1/2/2b coding AR-V3, pCMV5-1/2/3b coding AR-V4, and pCMV5-v567es coding AR-V567ES/AR-V12) were provided by Scott Dehm (The University of Minnesota) [43]. pCMV5-hAR encoding full-length AR (FL-AR) was acquired from Addgene (plasmid #89078). The pSV-S515A-AR, pSV-S515E-AR, pSV-S515D-AR plasmids were synthesised from the pSV-AR template using the QuikChange Lightning Multi Site-directed mutagenesis kit (Agilent Technologies). Mutations were introduced using the following primers: 5’-GCAGAGTGCCCTATCCC<u>GCT</u>CCCACTTGTGTCAAAA-3’ for pSV-S515A-AR, 5’-GCAGAGTGCCCTATCCC<u>GAG</u>CCCACTTGTGTCAAAA-3’ for pSV-S515E-AR, and 5’-GCAGAGTGCCCTATCCC<u>GAT</u>CCCACTTGTGTCAAAA-3’ for pSV-S515D-AR. Competent DH5α E. coli cells were transformed with the plasmids, plasmid DNAs were purified using QIAprep Spin Miniprep Kit (QIAGEN) and subjected to Sanger sequencing (GeneWiz) to confirm the presence of the expected mutations.

### Calcium phosphate-mediated transfections

COS-1 cells were transfected as previously described [42] with 1μg luciferase reporter (TAT-GRE-E1B-LUC), 50ng AR expression vector or empty vector and 50ng BOS-β-galactosidase per well. Cells were incubated for 24□hours in medium containing drug treatments.

### Luciferase and β-galactosidase assays

LNCaP/Luc or transfected COS-1 cells were lysed with Reporter Lysis Buffer (Promega). Luciferase activity was assayed using the Steadylite Plus kit (PerkinElmer). β-galactosidase activity was assayed using the Galacto-Light Plus system (Invitrogen). Light emission was measured using a VICTOR Light luminometer (PerkinElmer). For LNCaP/Luc lysates, luciferase activity was normalised to protein concentration. For COS-1 lysates, luciferase activity was normalised to β-galactosidase activity.

### C4-2B human CRPC xenografts

C4-2B cells were expanded by passaging twice weekly and harvested during the exponential growth phase. Cells were counted and viability was assessed using the trypan blue exclusion assay before implantation. Male NSG mice (6-8-week-old) purchased from Charles River (UK) were allowed to acclimatise for 1 week. Mice were injected subcutaneously with 2.5×10^6^ C4-2B cells resuspended in a volume of 100μL serum-free media and Matrigel Basement Membrane Matrix High Concentration (Corning) in a 1:1 ratio. Animals were randomised into treatment groups when tumour volume reached 90mm^3^. Mice were treated at 4mL/kg body weight with vehicle (5% DMSO and 30% SBE-β-CD dissolved in distilled water), 50mg/kg CT7001, 25mg/kg enzalutamide, or 50+25mg/kg CT7001+enzalutamide (combination). Treatments were administered orally once daily for 21 days. Tumour size was measured every 3-4 days using digital calipers, and tumour volumes were calculated using the formula: length*width*height*π/6. Mice were sacrificed on the last day of treatment, 2 hours after the administration of the final dose.

### Mouse plasma assays

Whole blood was collected via cardiac puncture in EDTA-coated tubes and spun immediately at 7500rpm for 5 minutes in a benchtop centrifuge. Plasma was collected, snap frozen in liquid nitrogen and sent to the Cambridge Biochemistry Assay Lab (Cambridge University Hospital) for analysis. Mouse aspartate transaminase (AST), mouse urea, and human free PSA levels were measured using a Siemens Dimension EXL analyser.

### Immunohistochemistry

Dissected tumours were fixed in 4% paraformaldehyde for 48 hours and embedded in paraffin blocks. Sections of 4μm were stained with the following antibodies: Ki67 (12202), P-S5 PolII (ab193467), and P-T161/T160 CDK1/CDK2 (ab201008). Staining was visualised using Dako REAL DAB+ Chromogen (Agilent Technologies). Slides were counterstained with haematoxylin. Staining was quantified in a blinded fashion in 2 different fields of view per tumour in 3 animals per group.

### Statistical analyses

Statistical analyses were carried out using GraphPad Prism v9.0. Pairwise comparisons were performed using the student’s t-test. Multiple comparisons were carried out using one-way or two-way ANOVA analysis followed by Dunnett’s or Šídák’s multiple comparisons test unless otherwise stated. *In vivo* tumour growth rate was defined as the slope of the linear regression described by tumour volume datapoints over time. All experiments were conducted with at least two biological repeats.

### RNA-sequencing and analysis

Total RNA was extracted from 30mg frozen tumour using the Monarch Total RNA Miniprep Kit (NEB). RNA library preparation and sequencing was performed by Novogene (UK) using 1μg total RNA. Briefly, sequencing libraries were generated using NEBNext Ultra RNA Library Prep Kit for Illumina (NEB). Library quality was assessed using the Ilumina Bioanalyser 2100 system (Agilent Technologies). The library preparations were sequenced on a NovaSeq 6000 System (Illumina) and paired-end reads were generated. Clean reads were obtained from raw reads in FASTQ format by removing adapters, poly-N sequences, and low-quality reads. Paired-end clean reads were aligned to the reference genome (GRCh38) using STAR v2.5 software. HTSeq v0.6.1 was used to count the read numbers mapped of each gene. Raw sequencing data and a matrix containing raw counts were submitted to the GEO repository (accession number GSE198488).

### Differential gene expression analysis

Differential expression analysis was performed using the DESeq2 R package (v1.28.1) [44]. Statistical significance was computed using the likelihood ratio test and the resulting P-values were adjusted using the Benjamini and Hochberg’s approach. Genes with an adjusted P-value (padj)<0.05 were assigned as differentially expressed. Gene clustering was performed using the degPatterns clustering tool from the DEGreport package (v1.24.1) [45].

### Gene set enrichment analysis

Gene set enrichment analysis (GSEA) was performed using GSEA 4.1.0 software (Broad Institute) using the normalised counts table estimated by DESeq2 for all genes and the Hallmark collection of gene sets (MSigDB) [46]. Statistical significance was computed using 10000 gene set permutations and the analysis was run using default parameters. The gene sets with false discovery rate (FDR)<0.1 were considered statistically significantly enriched.

## RESULTS

### CT7001 target engagement in LNCaP prostate cells

Previous *in vitro* kinase assays showed CT7001 selectively targets CDK7, although higher concentrations additionally inhibited the activities of other CDKs [30]. Transitioning from biochemical to cellular environments may alter target activity and selectivity (e.g. due to differences in protein structure and accessibility or due to off-target binding). Therefore, we carried out a cellular thermal shift assay (CeTSA) in intact LNCaP cells to investigate which CDK targets are engaged by CT7001 in a cellular model with relevance to prostate cancer. In this assay, binding of a drug to a target leads to formation of a drug-target complex with shifted heat-stability relative to the unbound target; as a result, the amount of soluble target increases in a dose-dependent manner following heat shock and can be determined by immunoblotting [37, 47].

To determine the melting characteristics for putative human CDK targets of CT7001, thermal proteome stability data for human cell lines were downloaded from the Meltome atlas [36]. The thermal profiles of CDK1, -2, -4, -7, and -9 were used to interpolate melting temperatures, T_m_, defined as the temperature which aggregates 50% soluble protein (**Figure 1A-B**). The T_m_ of the different CDKs ranged between ∼47-53°C. The Meltome data for CDK7 were validated in LNCaP cells (**Figure 1C(i)**). We next confirmed that heating of live LNCaP prostate cells at 54°C for 3 minutes decreases soluble CDK levels sufficiently to allow exploration of target engagement within the same cell suspension (**Figure 1C(ii)**). CeTSA isothermal dose-response fingerprints were then generated at 54°C using intact LNCaP cells treated with increasing concentrations of CT7001 (0-20μM) for 3 hours. Stabilization of CDK7, CDK2, and CDK9, but not of CDK4 or CDK1, was observed, as illustrated by increased protein levels compared with DMSO treatment (**Figure 1D(i)**). Additionally, we noted earlier engagement with CDK7, consistent with the compound’s reported selectivity (**Figure 1D(ii)**) [30]. Overall, these data suggest that CT7001 can bind preferentially to CDK7 but has a degree of engagement also with CDK2 and CDK9 at higher concentrations, which could contribute to efficacy in preclinical models.

**Figure 1.**
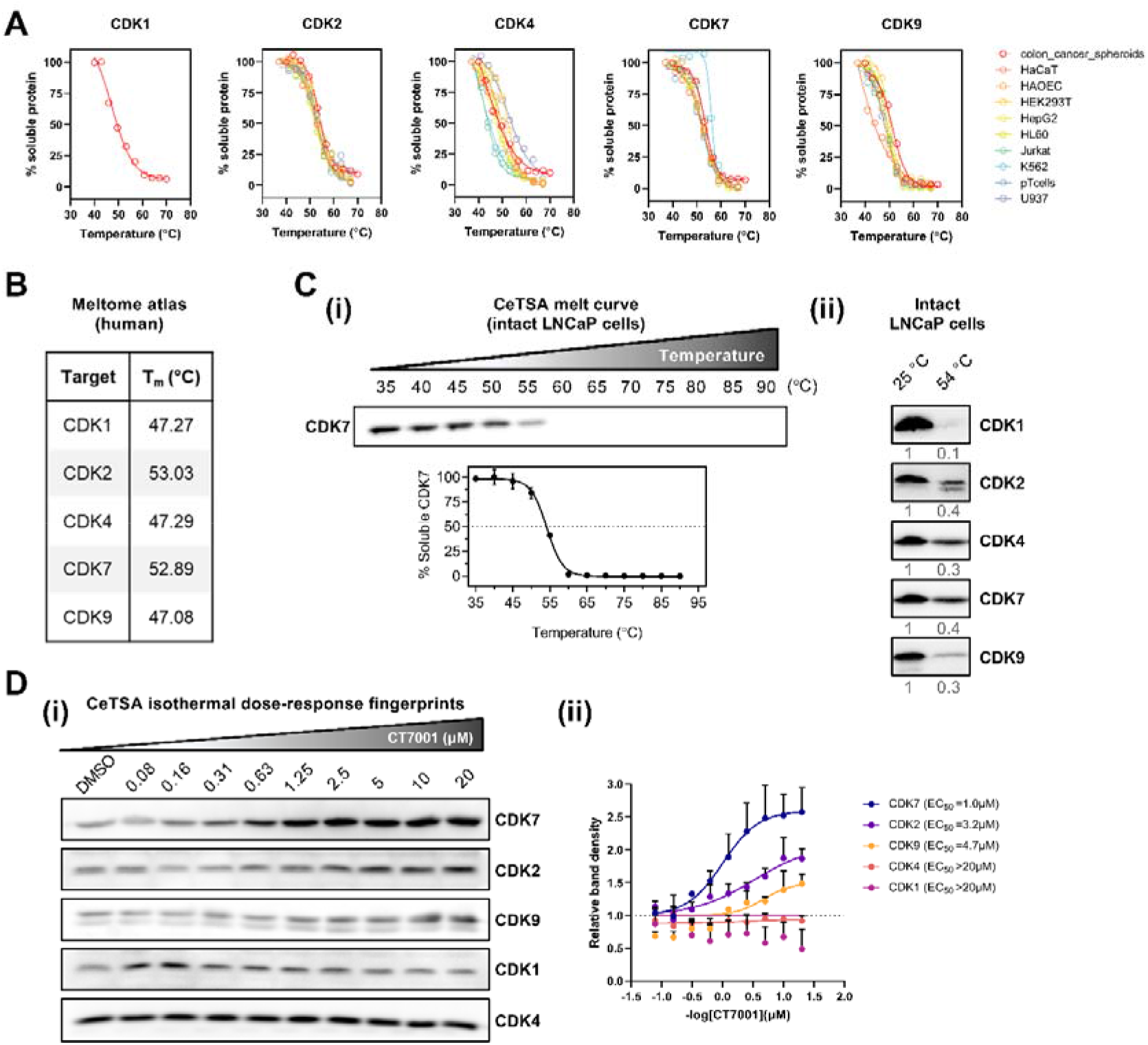
CT7001 target engagement in live LNCaP prostate cells. (A) Thermal stability profiles of putative human CDK targets, using data from the Meltome atlas. (B) Calculated melting temperatures (T_m_) of human CDK targets. (C) (i) Validation of the CDK7 melt curve in live LNCaP cells and (ii) immunoblots showing the effect of heat shock (54°C) on soluble levels of CDK1, CDK2, CDK4, CDK7, and CDK9. Numbers underneath blots represent remaining fraction of soluble protein. (D) (i) Cellular thermal shift assay (CeTSA) isothermal dose-response fingerprints at 54°C in intact LNCaP cells treated with the indicated CT7001 concentrations for 3 hours and (ii) quantification of target engagement relative to untreated samples. All immunoblots are representative of 3 biologically independent experiments.

### CT7001 inhibits cell growth and promotes cell cycle arrest *in vitro*

The ability of CT7001 to inhibit growth of prostate cancer cell lines was evaluated using growth-rate (GR) inhibition studies. GR curves and associated GR metrics are not confounded by the number of cell divisions taking place over the course of the experiment (as opposed to percent viability curves and traditional drug sensitivity metrics) [40], thus representing an improved methodology to measure drug sensitivity and have been proposed as replacements for conventional metrics [48]. Dose-dependent GR inhibition was observed in all cell lines treated with CT7001 over 72 hours (**Supplementary Figure S1A**). We calculated the drug concentration that reduces GR by 50% (GR_50_, the primary GR metric for drug potency) as well as the maximal measured GR value (GR_Max_, the primary GR metric for drug efficacy), and summarised these values for each prostate line tested in **Figure 2A**. GR_50_ ranged between 0.08-0.65μM for malignant prostate cell lines, suggesting potent growth inhibition, while negative GR_Max_ values were indicative of cytotoxicity in all lines except PC3 and DU145. GR_50_ concentrations for the two non-malignant cell lines (BPH-1 and PNT1A) were respectively 3.5-fold and 1.7-fold higher than the average GR_50_ for malignant lines, suggestive of lower drug potency in normal/benign prostate cells. In LNCaP cells, inhibition of CDK7 activity was associated with markedly reduced retinoblastoma (Rb) phosphorylation, a downstream target of cell cycle CDKs, while levels of CDK7, cyclin H and MAT1 were unaffected (**Figure 2B**). In contrast, decreased PolII CTD phosphorylation at S5 (a direct substrate of CDK7) and at S2 (phosphorylation mediated by CDK9) was only observed at concentrations above the GR_50_ of prostate cell lines and was temporary for S5 (**Figure 2C**). This indicates that therapeutic concentrations of CT7001 are sufficient to suppress the Rb-pathway but are unlikely to cause global PolII transcription inhibition in prostate cancer cells. Cell cycle analysis of asynchronous cell populations treated with CT7001 and stained with propidium iodide showed significant reduction of cells in S-phase, concomitantly with an increase in diploid (G_0_/G_1_) cells in LNCaP, C4-2B, and, to a lesser extent, in DU145 cells, consistent with Rb inhibition, and an increase in tetraploid (G_2_/M) cells in PC3 cells (**Figure 2B and Supplementary Figure S1B**). Mitosis markers (phosphorylated Rb, CDK1/CDK2, and histone H3) were decreased in LNCaP and PC3 cells, confirming a reduction in the fraction of proliferative cells (**Supplementary Figure S1C**). Collectively, these data support that prostate cancer cell growth and cell cycle progression are substantially disrupted by CT7001 treatment.

**Figure 2.**
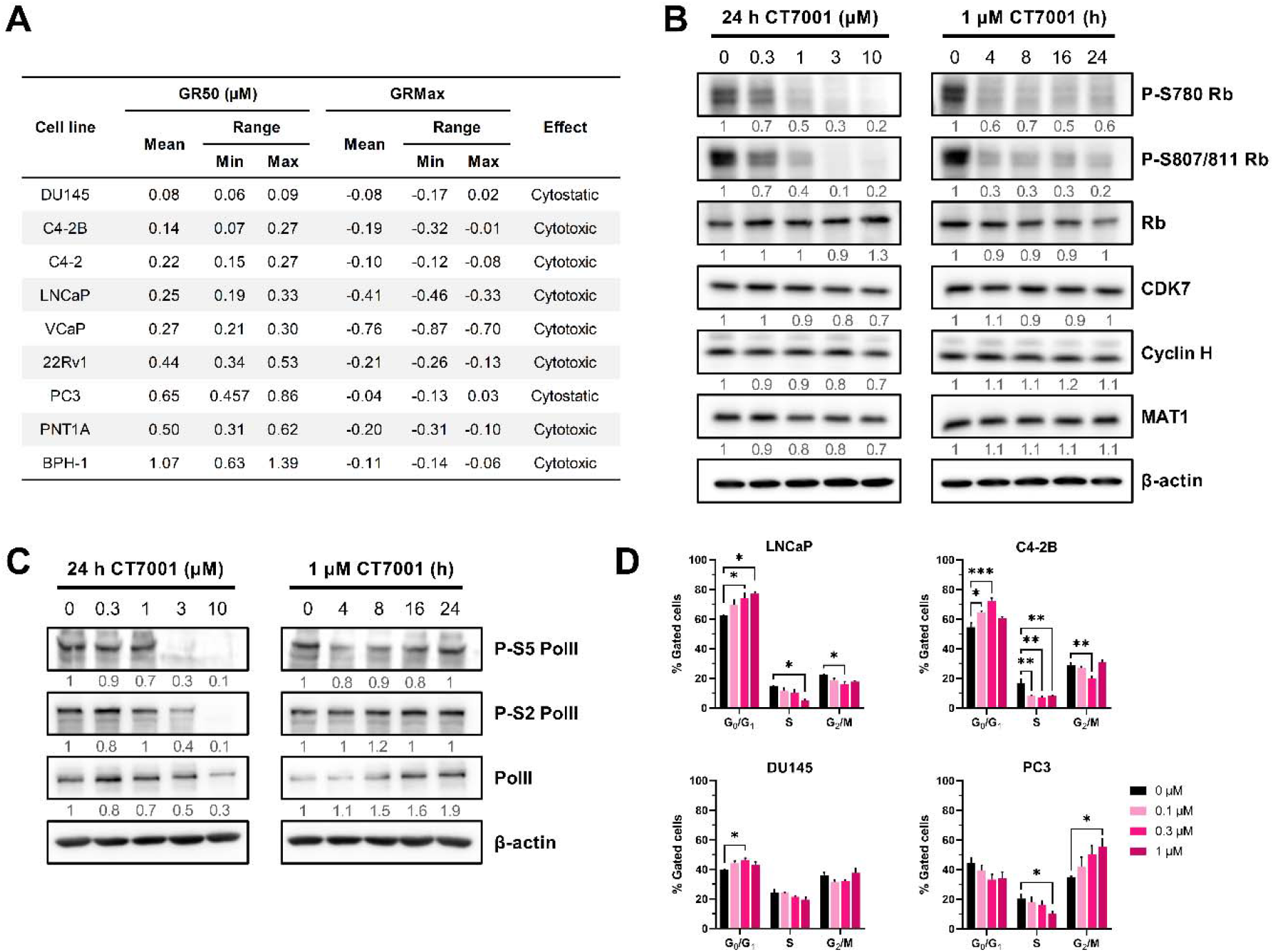
CT7001 inhibits proliferation of prostate cell lines and disrupts cell cycle progression. (A) Table summarising growth-rate (GR) metrics in prostate lines treated with CT7001 for 72 hours (n=3-4 per cell line). (B) Immunoblots from LNCaP cells treated with CT7001 showing effect on Rb phosphorylation and CAK expression (representative of n=3). (C) Immunoblots from LNCaP cells treated with CT7001 showing effect on PolII phosphorylation (representative of n=3). Numbers underneath blots represent relative band density quantified across 3 independent repeats. Data were adjusted to the loading control (β-actin). (D) FACS analysis of prostate cancer cell lines treated with CT7001 for 72 hours and stained with propidium iodide showing cell cycle distributions (n=3). Data are presented as mean ± SEM. P-values were determined using one-way ANOVA followed by Dunnett’s multiple comparisons test.

### CT7001 triggers activation of the p53 tumour suppressor pathway

A cytotoxic tendency was observed in GR studies in most prostate lines except DU145 and PC3 cells, which are p53-deficient. This prompted us to investigate whether CT7001 causes activation of the p53 pathway. Indeed, accumulation of p53 protein and increased serine-15 (S15) phosphorylation occurred in p53-intact LNCaP and C4-2B cells after 48 hours of treatment, but not in loss-of-function p53-mutant DU145 cells or PC3 cells (**Figure 3A**). Activation of p53 was most obvious following treatment with CT7001 concentrations over 1μM and was concomitant with increased levels of cleaved PARP1, a marker of apoptosis, and of p21, a CDK inhibitor which promotes cell cycle arrest. Further, transcriptional activation of known p53 target genes was confirmed in LNCaP cells but not in PC3 or DU145 cells (**Figure 3B**) and caspase 3/7 assays confirmed a significant increase in apoptosis in LNCaP cells, but not in PC3 cells, upon treatment with CT7001 concentrations up to 10μM (**Figure 3C**). These data indicate activation of p53 signalling contributes to cell cycle arrest and apoptosis in response to CT7001 treatment.

**Figure 3.**
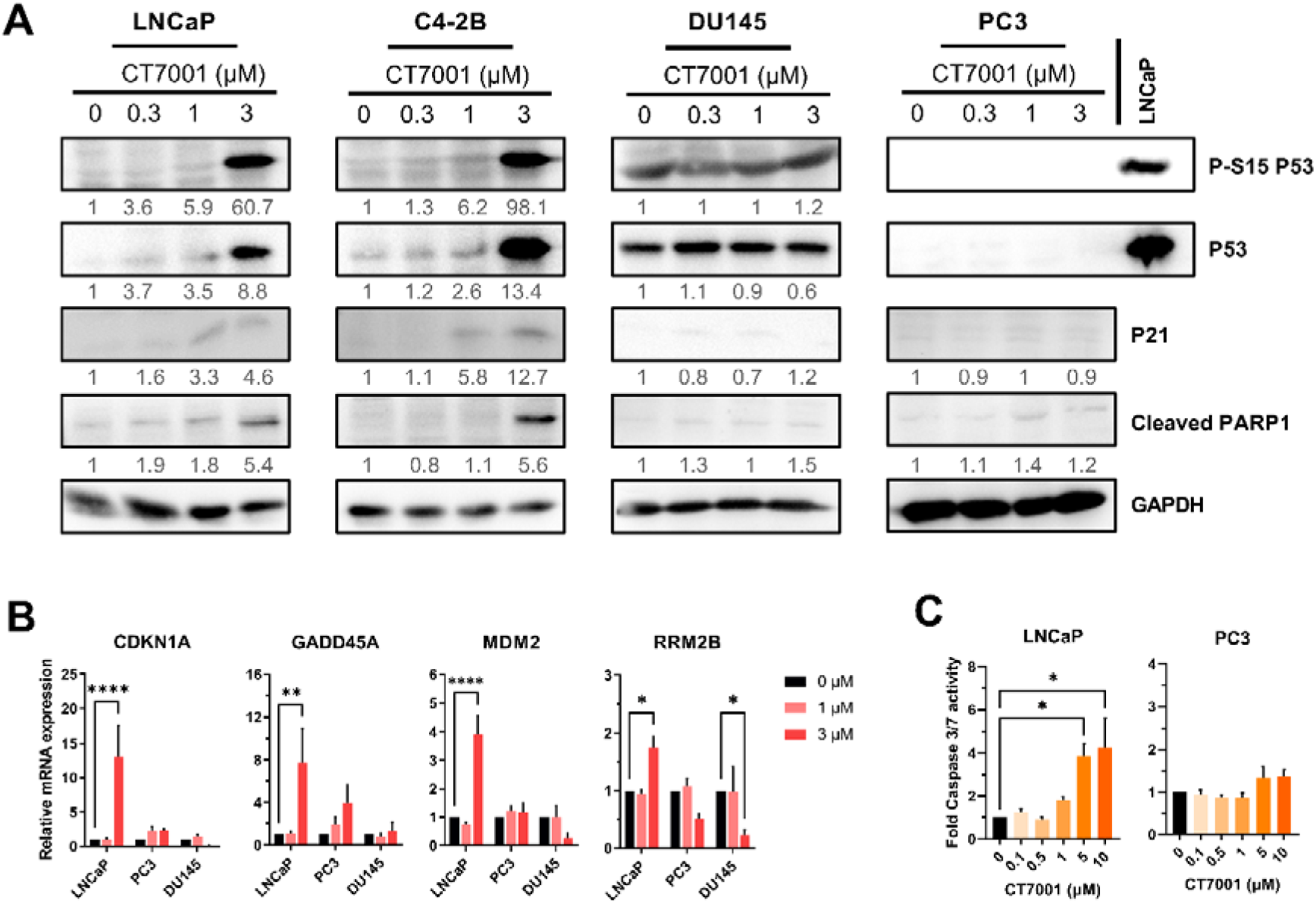
CT7001 induces activation of the p53 tumour suppressor and promotes apoptosis. (A) Immunoblots from prostate cancer cell lines treated with CT7001 for 48 hours (representative of n=3). Numbers underneath blots represent relative band density quantified across 3 independent repeats. Data were adjusted to the loading control (GAPDH). For PC3 blot, LNCaP extract was included as a positive control. (B) RT-qPCR analysis of p53 gene targets in LNCaP, PC3, and DU145 cells treated with CT7001 for 24 hours (n=3). P-values were determined using two-way ANOVA followed by Šídák’s multiple comparisons test. (C) Induction of apoptosis monitored via caspase 3/7 assays in LNCaP and PC3 cells following treatment with CT7001 for 72 hours (n=3). P-values were determined using one-way ANOVA followed by Dunnett’s multiple comparisons test. Data are presented as mean ± SEM.

### CT7001 impairs androgen-induced transactivation of AR *in vitro*

Given previous reports suggesting a coactivator role of CDK7 in AR-dependent transcription [23, 25], we sought to explore the ability of CT7001 to inhibit AR transactivation. We used the LNCaP/Luc cell line, which has endogenous expression of an active mutant (T877A) form of AR, and an integrated androgen-responsive luciferase reporter [35]. Androgen treatment (mibolerone) induced AR reporter activity, which was suppressed in a concentration-dependent manner by CT7001 (**Figure 4A**). Furthermore, mRNA expression of the endogenous AR target gene PSA also decreased in response to CT7001 treatment (**Figure 4B**). This effect was not due to reduced AR levels (**Supplementary Figure S2A**), impaired AR nuclear translocation in response to ligand (**Supplementary Figure S2B-C**), or reduced AR chromatin binding at well-characterised androgen response elements (**Supplementary Figure S2D**).

**Figure 4.**
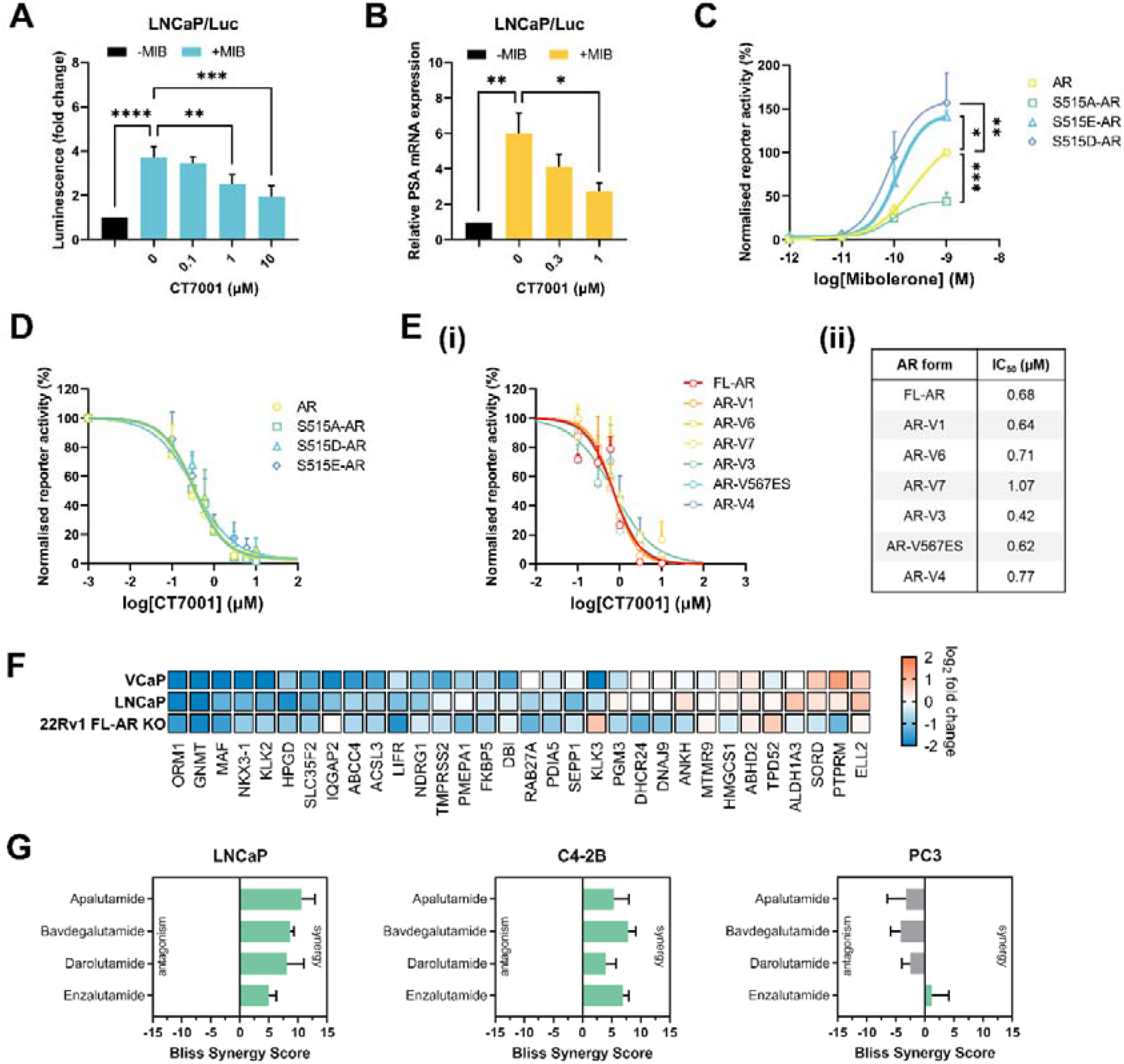
CT7001 interferes with AR transactivation. (A) Luciferase assay in LNCaP/Luc cells treated with CT7001 in the presence of 1nM mibolerone (MIB) for 24 hours (n=6). Data are presented as fold change relative to no androgen control. (B) PSA mRNA expression in LNCaP/Luc cells treated as in (A) (n=3). (C) AR reporter activity in COS-1 cells transfected with wild-type or S515 mutant AR plasmids. Cells were treated for 24 hours. Responses were normalised to β-galactosidase for transfection efficiency. Data are presented as % activity relative to wild-type AR, with the highest response representing 100% activity (n=4-5). (D) AR reporter activity in COS-1 cells transfected as in (C) and treated with CT7001 for 24 hours in the presence of 1nM MIB (n=3-4). (E) (i) AR reporter activity in COS-1 cells transfected with full-length AR (FL-AR) or AR variants (AR-V) plasmids. Transcriptional repression by CT7001 was assessed after 24 hours in the presence of 1nM MIB (n=3-4). (ii) Table summarising AR reporter activity IC_50_ for CT7001 treatment of FL-AR and AR-V. (F) Gene expression (RT-qPCR) of 32 AR targets in prostate lines. Cells in presence of androgen were treated ±1μM CT7001 for 24 hours, the colour represents fold change in presence vs absence of CT7001 (n=2). (G) Bliss synergy scores for CT7001 combinations with AR-targeting compounds (n=4). Data are presented as mean ± SEM. P-values in A-C were determined using one-way ANOVA followed by Dunnett’s multiple comparisons test.

To investigate whether the mechanism of AR repression involves the reported phosphorylation of AR by CDK7 at S515, we employed AR luciferase reporter assays in COS-1 cells transfected with expression vector encoding either wild-type AR or AR in which S515 was replaced either by non-phosphorylatable alanine (S515A-AR), or by the phosphomimetic aspartic acid (S515D-AR) or glutamic acid (S515E-AR).

Cells were co-transfected with an androgen-responsive luciferase reporter and β-galactosidase reporter as internal transfection efficiency control. Androgen treatment stimulated the transcriptional activity of wild-type AR and mutant AR proteins in a concentration-dependent manner (**Figure 4C**). Relative transcriptional activity was lower than wild-type AR for the S515A-AR mutant and greater for the S515E-AR and S515D-AR mutants, consistent with previous reports [24, 25, 49]. We reasoned that if decreased phosphorylation of AR at S515 is essential for the inhibitory activity of CT7001, then the S515E and S515D mutations would abrogate its effect. However, treatment with CT7001 suppressed transactivation of wild-type and S515 AR mutants with similar potency and efficacy (**Figure 4D**), suggesting CT7001 suppresses AR-driven transcription via a different mechanism.

We next sought to investigate whether AR repression in response to CT7001 treatment requires a functional AR LDB. Similar reporter assays were carried out in COS-1 cells using plasmids encoding full-length AR (FL-AR) or constitutively active AR-Vs that lack the LBD (depicted in **Supplementary Figure S2E**) which are insensitive to androgen treatment (**Supplementary Figure S2F**), and for which there are no approved inhibitory compounds. Treatment with CT7001 repressed transcription mediated by FL-AR with EC_50_ of 0.68μM, and by all AR-Vs tested, with EC_50_ ranging between 0.42-1.07μM (**Figure 4E**). To validate our findings from AR luciferase reporter assays and investigate transcriptional effects in further prostate cancer cell lines, we measured transcript levels of 32 known AR target genes, comprising an AR activity signature (a list of TaqMan qPCR assays is provided in **Additional File 2, Supplementary Table 2**). In LNCaP cells, which express FL-AR, and VCaP cells, which express both FL-AR and AR-V, treatment with 1μM CT7001 repressed expression of most AR targets (**Figure 4F**). We repeated this experiment in the 22Rv1 FL-AR KO cell line, which completely lacks FL-AR and in which expression of AR targets is maintained solely by constitutively active AR-Vs [33]. As in the other cell lines, treatment with 1μM CT7001 was sufficient to decrease the expression of most AR target genes measured (**Figure 4F**). The results from AR reporter assays and gene expression analyses of AR target genes in prostate cancer cell lines support the hypothesis that CT7001 represses transcription mediated by both full-length AR and truncated, constitutively active AR-Vs *in vitro*. Collectively, these indicate AR pathway suppression as a secondary mode of action of CT7001 in prostate cancer cells, in addition to cell cycle inhibition. Thus, we next explored potential synergic effects on cell growth between CT7001 and four AR-targeted compounds: the second-generation antiandrogens apalutamide, darolutamide, and enzalutamide, and the AR-specific protein degrader bavdegalutamide/ARV-110. Co-treatment with CT7001 and AR-targeted therapies resulted in additive-to-synergistic growth inhibition in the AR-positive hormone-dependent LNCaP and CRPC C4-2B cell lines, as indicated by positive Bliss independence scores (**Figure 4G** and **Supplementary Figure S3**). The combinations were, as expected, not synergistic in the AR-negative CRPC PC3 cell line.

### CT7001 and enzalutamide supress tumour growth *in vivo* in an additive manner

To assess tumour growth inhibition of CT7001 alone or in combination with enzalutamide, a widely used antiandrogen, immunocompromised (NSG) male mice with established subcutaneous C4-2B xenografts were assigned into 4 treatment groups: vehicle, CT7001 (50mg/kg) alone, enzalutamide (25mg/kg) alone, or a combination of enzalutamide and CT7001. On day 21 of daily oral treatment, CT7001 and enzalutamide monotherapy reduced final tumour volume by 52% and 50% respectively, while combination therapy reduced final tumour volume by 73%, compared to vehicle treatment (**Figure 5A**). A significant reduction in the weights of excised tumours was also evident for all treatments (**Figure 5B**). Regression analysis using longitudinal tumour volume data showed a significant reduction in growth-rate in all treatment groups compared to vehicle (vehicle vs. CT7001 p=0.0006; vehicle vs. enzalutamide p<0.0001; vehicle vs. combination p<0.0001) and, additionally, a significant reduction in tumour growth-rate in the combination arm compared to enzalutamide alone (p=0.0156) (**Figure 5C**). Mean tumour doubling time was 8 days for vehicle-treated mice, 12 days for CT7001-treated mice, 11 days for enzalutamide-treated mice, and 21 days for mice treated with both CT7001 and enzalutamide (**Figure 5D**). Tumour growth inhibition was also associated with significantly reduced plasma free PSA in enzalutamide- and combination-treated mice relative to vehicle controls, and with a similar trend in CT7001-treated mice (**Figure 5E**). All treatments were generally well-tolerated, with less than 5% body weight loss on average in all treatment arms, which was comparable to the weight loss observed in vehicle controls (**Figure 5F**). Additionally, plasma AST and urea levels appeared unchanged, except for a small reduction in plasma urea concentration in enzalutamide-treated mice relative to vehicle controls (**Supplementary Figure S4A-B**). There were no obvious histopathological changes in the liver of treated animals (**Supplementary Figure S4C**). Immunohistochemistry of resected tumours showed CT7001 monotherapy reduced Ki67 expression, with further decrease observed in tumours treated with a combination of enzalutamide and CT7001 (**Figure 5G-H**). In addition, T-loop phosphorylation of CDK1/CDK2 was additively decreased by CT7001 and enzalutamide treatments, and a small decrease in PolII phosphorylation at S5 was detected in tumours treated with CT7001 alone or in combination with enzalutamide.

**Figure 5.**
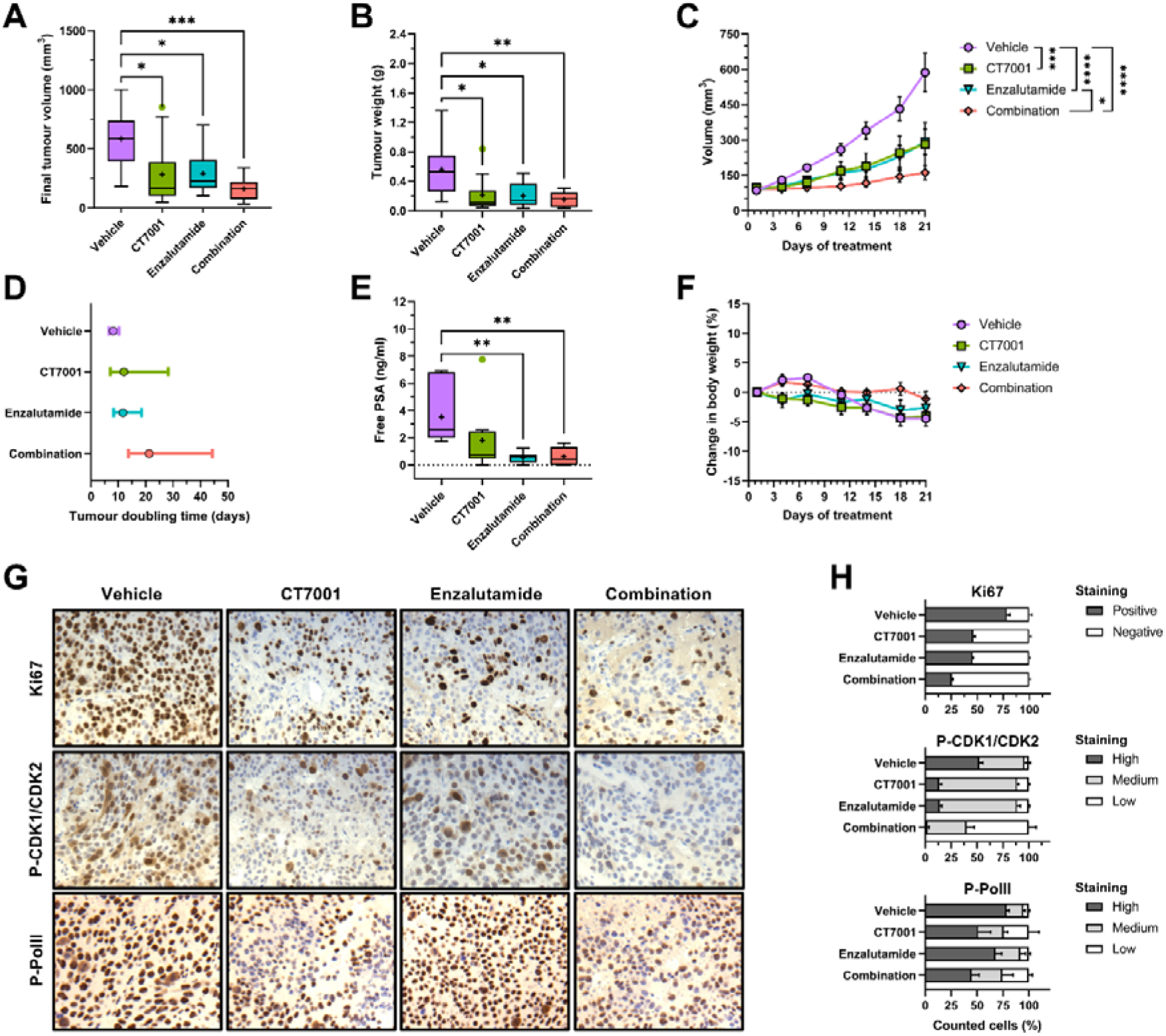
CT7001 administration prevents CRPC growth *in vivo* and has additive effects in combination with enzalutamide. (A) Tumour volumes on day 21 for male NSG mice bearing C4-2B xenografts following daily oral treatment with vehicle, CT7001 (50 mg/kg), enzalutamide (25 mg/kg) or combination (n=10 per group). (B) Resected tumour weights at the end of 21 days of treatment. (C) Mean tumour volumes ± SEM over the course of treatment. Linear regression showed statistically significant differences in growth between different treatment groups. (D) Tumour doubling times ± 95% confidence intervals. (E) Plasma free prostate specific antigen (PSA) levels in treated mice. (F) Mean change in body weight ± SEM of treated mice over the course of 21 days of treatment. (G) Images of Ki67, P-T161/T160 CDK1/CDK2 and P-S5 RNA polymerase II (PolII) immunohistochemistry staining of resected C4-2B xenografts. Images are representative for 3 animals per group. (H) Quantification of immunohistochemistry staining in n=3 animals per group (mean ± SEM). P-values were determined using one-way ANOVA followed by Šídák’s multiple comparisons test. Data in (A), (B) and (E) are presented using Tukey boxplots. Mean values are plotted as a “+” and outliers are shown as individual datapoints. *p<0.05; **p<0.01; ***p<0.001; ****p<0.0001.

Finally, we sought to identify drug-induced transcriptome changes and biological pathways associated with treatment response by carrying out RNA-seq analysis of the treated C4-2B xenografts (n=3-4 per group). Unsupervised principal component analysis (PCA) captures 28.94% of the gene expression variance in the first two PCs and shows separation of the different groups, with the enzalutamide and combination groups clustering furthest from the vehicle group (**Figure 6A**). We performed differential gene expression analysis using DESeq2 [44] and computed statistical significance using a likelihood ratio test. This analysis identified 1915 differentially expressed genes that showed significant changes in expression across the four different groups (padj<0.05). Clustering analysis performed using the DEGreport package [45] identified 6 major differential gene expression clusters (**Figure 6B-C**). Clusters 1-4, representing a large majority of 1612 genes, showed altered expression in response to enzalutamide and combination treatment, while treatment with CT7001 alone led to only small changes in expression. This pointed to enzalutamide as the major driver of transcriptional effects in response to combination therapy. We performed gene set enrichment analysis (GSEA) using a matrix containing DESeq2-normalised counts for all detectable genes in the different groups and calculated enrichment scores, relative to vehicle, for the Hallmark gene sets collection [46] (**Figure 6D**). With respect to CT7001 treatment alone, the results of the GSEA are consistent with the effects of CT7001 treatment observed *in vitro*. Negative enrichment for several cell cycle-related gene sets (e.g. G2M checkpoint, E2F targets, and mitotic spindle) and the hallmark androgen response, and positive enrichment for the hallmark p53 pathway were noted (**Supplementary Figures S5 and S6**). In tumours treated with enzalutamide alone or with a combination of enzalutamide and CT7001, GSEA identified negative enrichment of several gene sets, including the hallmark androgen response, MYC targets, reactive oxygen species and fatty acid metabolism (**Figure 6D**). Reassuringly, the top negatively regulated gene set for enzalutamide and combination treatments was the hallmark androgen response; interestingly, this was also the case for CT7001 alone. A few hallmark gene sets were exclusively negatively enriched in the combination group (e.g. oxidative phosphorylation, glycolysis, estrogen responses, DNA repair). However, the top gene sets enriched in the combination group substantially overlapped with those enriched in the enzalutamide group (**Figure 6E**). This indicates that addition of CT7001 to enzalutamide at therapeutic doses contributes little to the transcriptional effect of enzalutamide and suggests cell cycle inhibition by CT7001 as the primary mechanism underpinning the additional growth repression observed in response to combination therapy.

**Figure 6.**
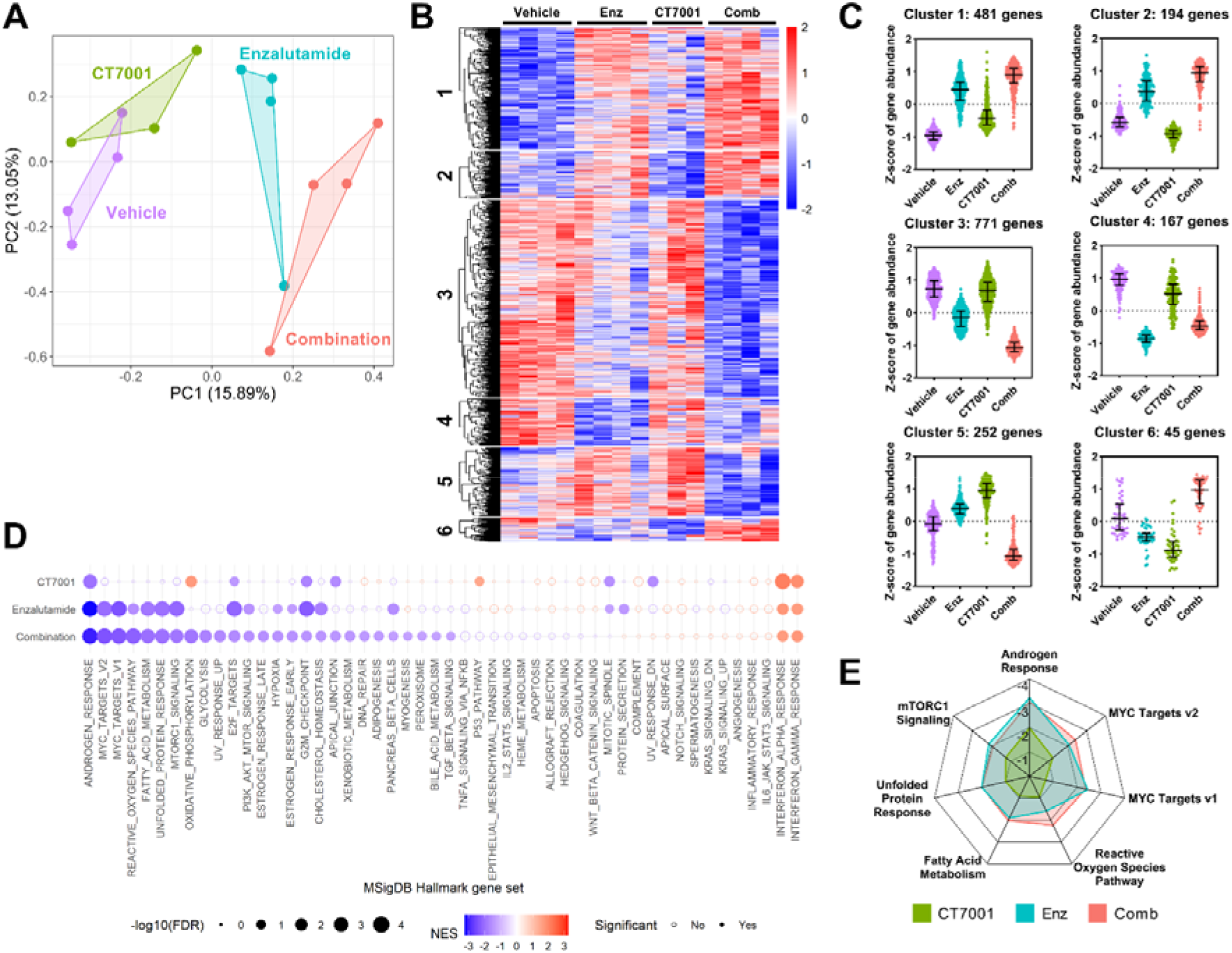
Transcriptomic analysis of C4-2B xenografts. (A) Unsupervised principal component analysis (PCA) clustering of treated xenografts. (B) Heatmap of differentially expressed genes identified through DESeq2 likelihood-ratio test as changing in expression across different treatments. (C) Clustering of differentially expressed genes based on expression profiles across different treatments. (D) Gene set enrichment analysis results showing MSigDB hallmark gene sets significantly enriched in treatment groups versus vehicle. Dot size represents -log_10_ false discovery rate (FDR) while colour indicates normalised enrichment score (NES). Significantly enriched gene sets (FDR<0.1) are shown using colour-filled dots. (E) Radar plot displaying normalised enrichment scores for CT7001, enzalutamide and combination for the top hallmarks gene sets enriched in the combination group.

## DISCUSSION

In this report, we present a systematic assessment of the mechanism of action and preclinical efficacy of CT7001 as a single agent or combined with the second-generation antiandrogen enzalutamide in established preclinical models of CRPC. Using cellular thermal shift assays in LNCaP cells, we show CT7001 preferentially targets CDK7 over CDK2 and CDK9. This complements published studies using cell-free *in vitro* kinase assays [30] and confirms CT7001’s potent and selective pharmacology against CDK7 in living prostate cancer cells. In contrast to one report suggesting CDK4 as a target of CT7001 [50], we found no evidence of direct engagement with CDK4 or with CDK1 at concentrations up to 20μM. Treatment with CT7001 induces potent growth inhibition and promotes cell cycle arrest in prostate cancer lines with submicromolar potency, while higher drug concentrations can induce activation of the p53-pathway and promote apoptosis. Our results show that CT7001 also inhibits the transcriptional activity of AR, a key transcription factor and oncogenic driver in prostate cancer. Increased sensitivity of AR-positive (compared to AR-negative) prostate cancer cell lines to THZ1, an inhibitor of CDK7/CDK12/CDK13, has been previously reported [23]. With respect to CT7001, potent growth inhibition was achieved in both AR-positive (LNCaP, C4-2, C4-2B, VCaP and 22Rv1) and AR-negative (DU145, PC3) lines.

Notwithstanding, complete loss of AR expression is rare in CRPC, where reactivation of AR signalling drives resistance to androgen deprivation therapy. This frequently involves mutation or loss of the AR LBD. In this report, we demonstrate that CT7001 treatment successfully inhibits growth and AR-dependent transcriptomes of cell lines modelling these scenarios: LNCaP cells carry a point mutation in the AR LBD (T877A) which confers promiscuous activation by alternative ligands, VCaP cells display AR gene amplification in addition to expressing constitutively active AR-Vs, while the engineered 22Rv1 FL-AR KO line displays ligand-independent AR transcriptome driven by AR-Vs and intrinsic resistance to AR targeting therapies. Direct inhibition of ectopically expressed AR-Vs was also demonstrated. As CT7001 mechanistically functions independently of the AR LBD, unlike all approved AR-targeted compounds, it is a good therapeutic candidate for CRPC tumours with AR reactivation. We also explored the therapeutic potential of combining CDK7 inhibition with the widely prescribed antiandrogen enzalutamide and found additive tumour growth repression *in vivo*, likely mediated largely through CT7001’s potent repression of cell cycle progression. Further research is warranted to determine whether, in models displaying resistance to antiandrogens, AR pathway suppression by CT7001 becomes essential for therapeutic efficacy.

Considering the more widespread effects of CDK7 on PolII-dependent transcription, mounting evidence suggests targeting transcriptional kinases using selective inhibitors could provide a sufficient therapeutic window to be effective without the initially feared toxicity [51, 52]. However, our data indicate that, while CT7001 treatment can reduce PolII phosphorylation *in vitro*, inhibition of the Rb and AR pathways is likely sufficient for therapeutic efficacy at low concentrations. This is in line with recent work using the CDK7-selective covalent inhibitor YKL-5-124, which caused no change in PolII phosphorylation at therapeutic doses [21, 22].

Several questions remain regarding the mechanism of action of CT7001 in prostate cancer. Firstly, activation of p53 presumably contributes to CT7001-induced apoptosis. The highly metastatic PC3 cell line, which does not express p53 protein, appears resistant to apoptosis in response to CT7001 treatment. In addition, activation of p53 transcriptional program has been demonstrated to sensitise some cancer cells to CDK7 inhibition by promoting pro-apoptotic pathways [53]. As the p53 tumour suppressor is frequently mutated in prostate cancer and has prognostic significance [4, 54], future studies should explore whether enhanced sensitivity or/and induction of apoptosis in response to CT7001 treatment is dependent on an intact p53-pathway. Indeed, preliminary data from the clinical trial strongly indicate that *TP53* status is associated with response to CT7001 therapy in patients with metastatic hormone receptor-positive/HER2-negative breast cancer [32]. Secondly, in addition to suppressing cell proliferation, CDK7 inhibition by YLK-5-124 was found to induce interferon gamma signalling along with other inflammatory response pathways in models of small cell lung cancer [22]. This, in turn, triggered robust immune cell signalling, which potentiated the antitumour immune response and enhanced response to anti-PD-1 immunotherapy. Whether CT7001 treatment can provoke similar antitumour immune responses in immunocompetent murine models of prostate cancer remains to be explored.

## CONCLUSIONS

In summary, the data presented here demonstrate that the orally bioavailable CDK7 inhibitor CT7001 impedes malignant cell growth by targeting proliferation pathways and, in addition, downregulates oncogenic AR signalling and induces apoptosis in CRPC models. This multifaceted mechanism of action drives additive combinatorial activity with the antiandrogen enzalutamide in the C4-2B xenograft model of advanced CRPC. Therefore, this study supports CDK7 inhibition as a therapeutic strategy for CRPC and provides a rationale for new combination regimens consisting of CDK7 inhibitors and antiandrogen therapy, all the more so as CT7001 demonstrated acceptable safety profile with evidence of target engagement in clinical trials.

## LIST OF ABBREVIATIONS

AR: androgen receptor
AR-V: androgen receptor variant
AST: aspartate aminotransferase
CAK: CDK-activating kinase
CDK: cyclin-dependent kinase
CeTSA: cellular thermal shift assay
CRPC: castration-resistant prostate cancer
CTD: C-terminal domain
DMSO: dimethylsulfoxide
FCS: foetal calf serum
FDR: false discovery rate
FL-AR: full-length androgen receptor
GR: growth-rate
GSEA: gene set enrichment analysis
LBD: ligand-binding domain
Padj: adjusted p-value
PBS: phosphate buffered saline
PCA: principal component analysis
PolII: RNA polymerase II
PSA: prostate-specific antigen
P-TEFb: positive transcription elongation factor b
Rb: retinoblastoma
RT-qPCR: quantitative reverse transcription PCR
SRB: sulforhodamine B

## DECLARATIONS

### Ethics approval

All mouse experiments were carried out by licensed investigators in accordance with the UK Home Office Guidance on the Operation of the Animal (Scientific Procedures) Act 1986 (HMSO, London, UK, 1990).

### Consent for publication

Not Applicable

### Availability of data and materials

The RNA-sequencing data generated in this study were deposited in NCBI Gene Expression Omnibus (GEO accession GSE198488). The data will be available following a 6-month embargo from the date of publication. The melting curves for CDK proteins were downloaded from the online Meltome Atlas (http://meltomeatlas.proteomics.wzw.tum.de:5003/). All other data generated or analysed during this study are included in this article and its supplementary information provided in **Additional file 1** and **Additional file 2**.

## Additional file 1.docx – Supplementary figures

Supplementary Figure S1. CT7001 treatment affects proliferation of prostate cell lines.

Supplementary Figure S2. Effect of CT7001 treatment on androgen receptor signalling.

Supplementary Figure S3. Bliss synergy matrices for CT7001 and AR-targeting therapies in prostate cancer cell lines.

Supplementary Figure S4. Additional plasma biochemistry in the mouse cohort following treatment and mouse liver haematoxylin & eosin staining.

Supplementary Figure S5. GSEA results for gene sets negatively enriched in CT7001 treated tumours.

Supplementary Figure S6. GSEA results for gene sets positively enriched in CT7001 treated tumours.

## Additional file 2.docx – Supplementary materials and methods

Supplementary materials and methods

Supplementary Table 1. Primer sequences for RT-qPCR.

Supplementary Table 2. TaqMan low-density array qPCR assays

Supplementary Table 3. Primer sequences for ChIP-qPCR.

Supplementary References

### Funding

T.A.C. was funded by the Imperial College Medical Research Council Doctoral Training Partnership and associated Supplement Scheme, and by the Imperial College London President’s PhD Scholarship scheme. We also acknowledge support from the Cancer Research UK Imperial Centre.

### Authors’ contributions

T.A.C., A.V.C., S. Ali, C.L.B. designed *in vitro* experiments. T.A.C., A.V.C., L.P., D.C. and C.L.B. designed *in vivo* experiments. T.A.C., A.V.C., K.K.G., S. Ang, L.P., and G.S.A. performed experiments and analysed data. A.O. conducted preliminary *in vitro* experiments underpinning this study. T.A.C., A.V.C., E.A., A.K.B., M.J.F., S. Ali, and C.L.B. contributed to data interpretation, drafting of the manuscript, and provided revisions. All authors read and approved the final manuscript.

## Acknowledgements

We thank Prof Ian Mills for providing the C4-2B cell line, Dr Luke Gaughan for providing the 22Rv1 and 22Rv1 FL-AR KO cell lines, Dr Scott Dehm for providing the pCMV5-CE1, pCMV5-CE2, pCMV5-CE3, pCMV5-1/2/2b, pCMV5-1/2/3b, and pCMV5-v567es plasmids, and Dr Chun Fui Lai for designing and providing the AR targets TaqMan arrays. We additionally acknowledge Joel Abrahams, Roxanne Wood, and the staff of the Imperial College Central Biomedical Services for their training and support in conducting *in vivo* xenograft studies, and Sophie Xue (Novogene, UK) for support conducting the xenograft RNA-sequencing project.

