## Supplemental Figures for "The CDK7 inhibitor CT7001 (Samuraciclib) targets proliferation pathways to inhibit advanced prostate cancer"

### SUPPLEMENTARY FIGURES

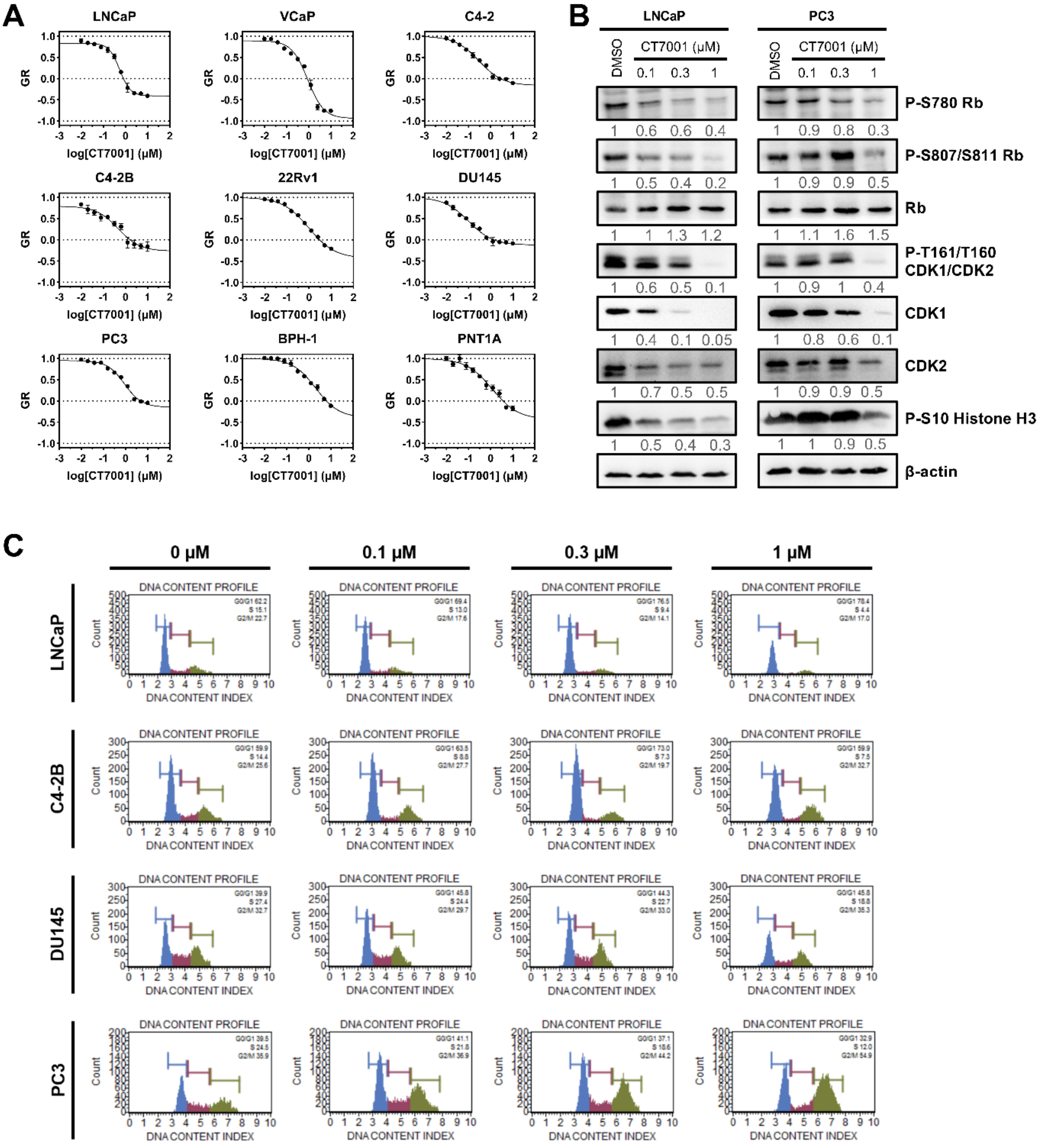


**Supplementary Figure S1. CT7001 treatment affects proliferation of prostate cell lines. (A)** Growth-rate (GR) inhibition curves in prostate lines treated with CT7001 for 72 hours. Relative cell numbers were measured using Sulforhodamine B assays (n=3-4 per cell line). **(B)** Representative immunoblots of LNCaP and PC3 cells treated with CT7001 for 72 hours showing decreased expression of proliferation markers (n=3). Numbers underneath blots represent relative band density quantified across 3 independent repeats. Data were adjusted to the loading control (β-actin). **(C)** DNA content profiles of asynchronous prostate cancer cell lines treated with CT7001 for 72 hours. Images are representative for 3 biologically independent experiments.

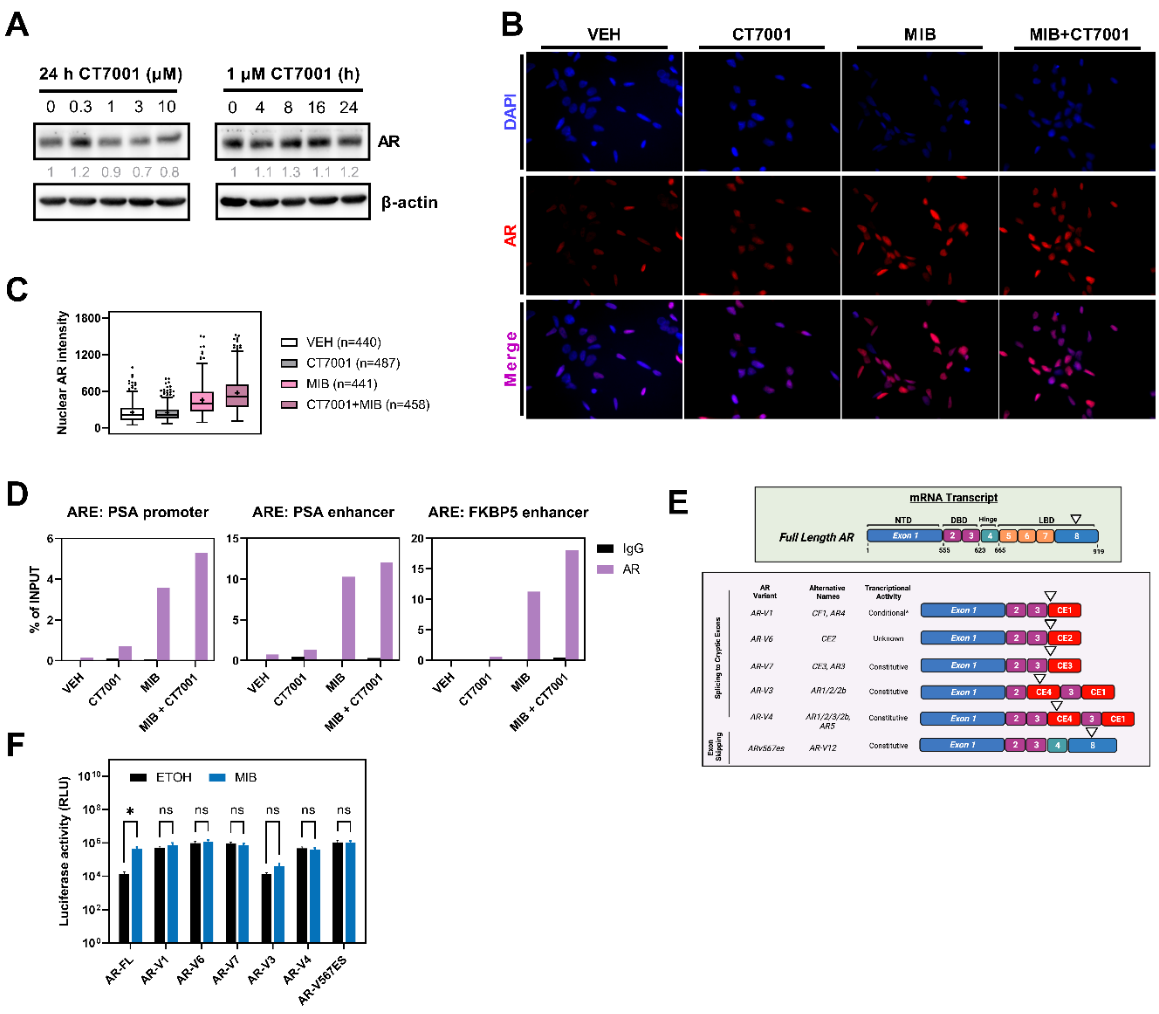


**Supplementary Figure S2. Effect of CT7001 treatment on androgen receptor signalling. (A)** Immunoblots of LNCaP cells treated with CT7001 as indicated. representative for 3 biologically independent experiments (representative of n=3). Numbers underneath blots represent relative band density quantified across 3 independent repeats. Data were adjusted to the loading control (β-actin). **(B)** Immunofluorescence images of LNCaP cells treated with vehicle or with 10µM CT7001 in the absence or presence of 10nM mibolerone (MIB) for 4 hours before fixation. Cells were stained with DAPI (blue) and with an anti-AR antibody (red) (representative of n=2). **(C)** Quantification of AR mean nuclear staining intensity for cells in (B). Tukey boxplots were used to present the data and mean values are plotted as a "+". Outliers are shown as individual datapoints. N represents nuclei quantified across 2 biologically independent experiments. **(D)** Chromatin-immunoprecipitation (ChIP)-qPCR in LNCaP cells treated with vehicle or with 10µM CT7001 in the absence or presence of 10nM mibolerone (MIB) for 4 hours before crosslinking. AR binding to androgen response elements (ARE) was quantified in the PSA gene promoter, PSA gene enhancer, and FKBP5 gene enhancer (representative of n=3). **(E)** Schematic diagram showing the mRNA transcript structure of full-length AR and truncated AR splice variants (AR-Vs). NTD=N-terminal domain; DBD=DNA-binding domain; LBD=ligand-binding domain; CE=cryptic exon. Created with BioRender.com. **(F)** AR reporter assays in COS-1 cells transfected with full-length AR (AR-FL) or truncated AR splice variants (AR-V) lacking the ligand binding domain. Cells were treated with vehicle (ethanol) or 1nM mibolerone (MIB) for 24 hours (n=4). Data are presented as mean ± SEM. P-values were determined using two-way ANOVA followed by Šídák's multiple comparisons test. ns=not significant; *p<0.05.

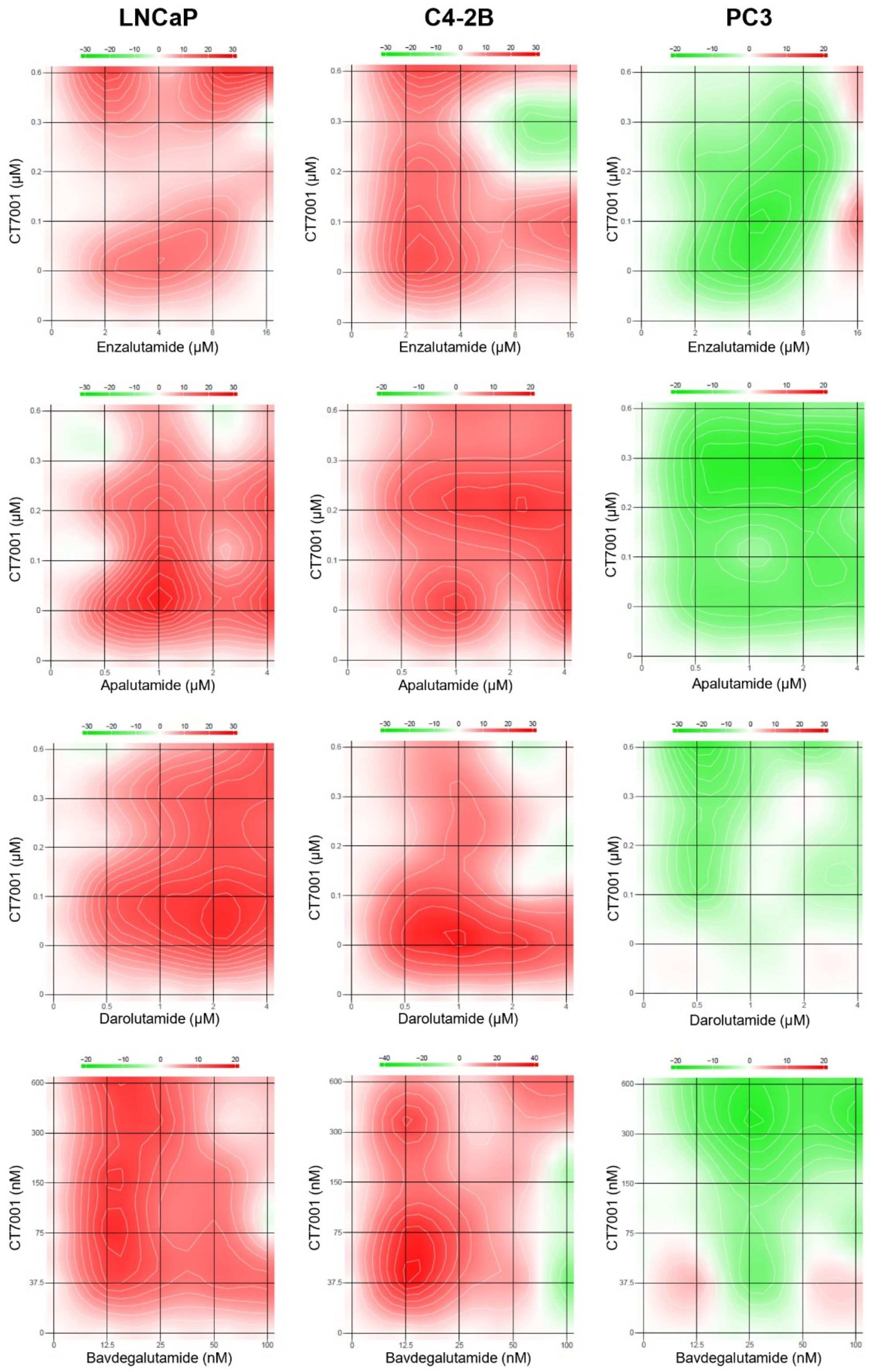


**Supplementary Figure S3. Bliss synergy matrices for CT7001 and AR-targeting therapies in prostate cancer cell lines.** Synergy values were determined using the Bliss reference model and the SynergyFinder 2.0 web application. Colour schemes represent Bliss independence scores. Red indicates synergistic interaction while green indicates antagonistic interaction (representative of n=4).

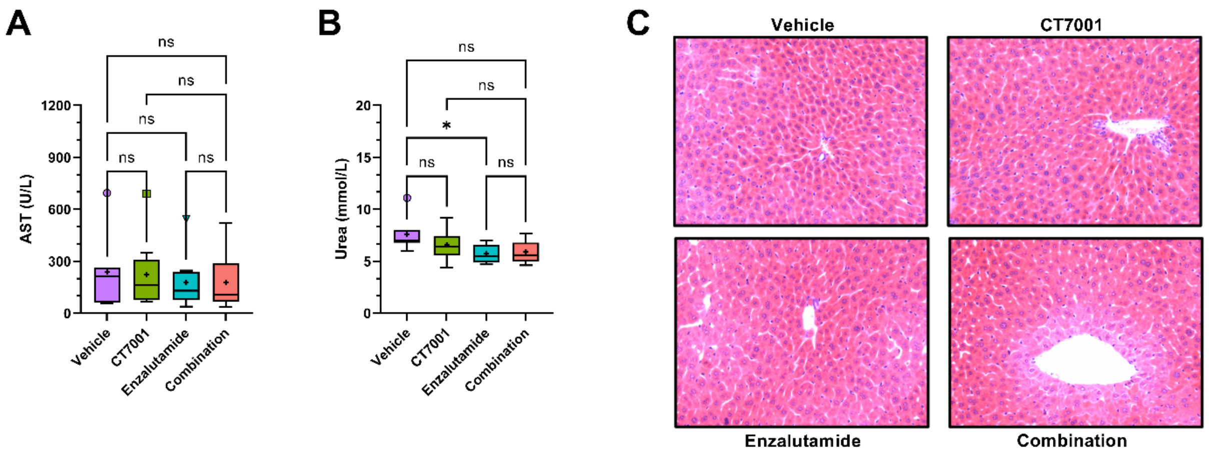


**Supplementary Figure S4. Additional plasma biochemistry in the mouse cohort following treatment and mouse liver haematoxylin & eosin staining. (A)** Measurement of plasma activity of aspartate transaminase (AST) from treated mice. **(B)** Measurement of plasma urea levels from treated mice. Tukey boxplots were used to present the data in (A) and (B). Mean values are plotted as a “+” and outliers are shown as individual datapoints. **(C)** Haematoxylin & eosin staining of liver sections from treated NSG mice (representative for n=3 animals per group).

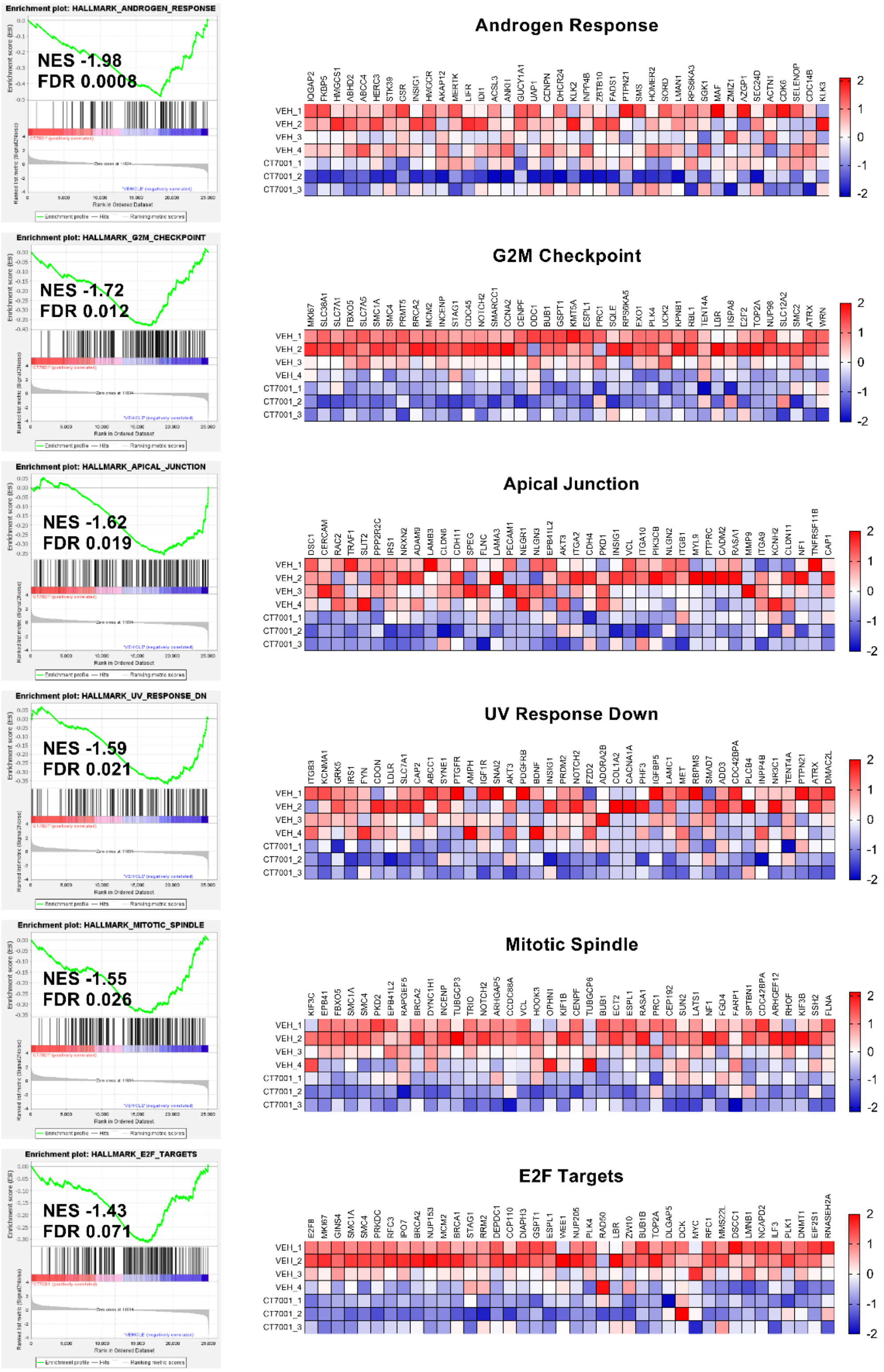


**Supplementary Figure S5. GSEA results for gene sets negatively enriched in CT7001 treated tumours.** Enrichment plots and leading edge heatmaps (top 40 genes) are shown.

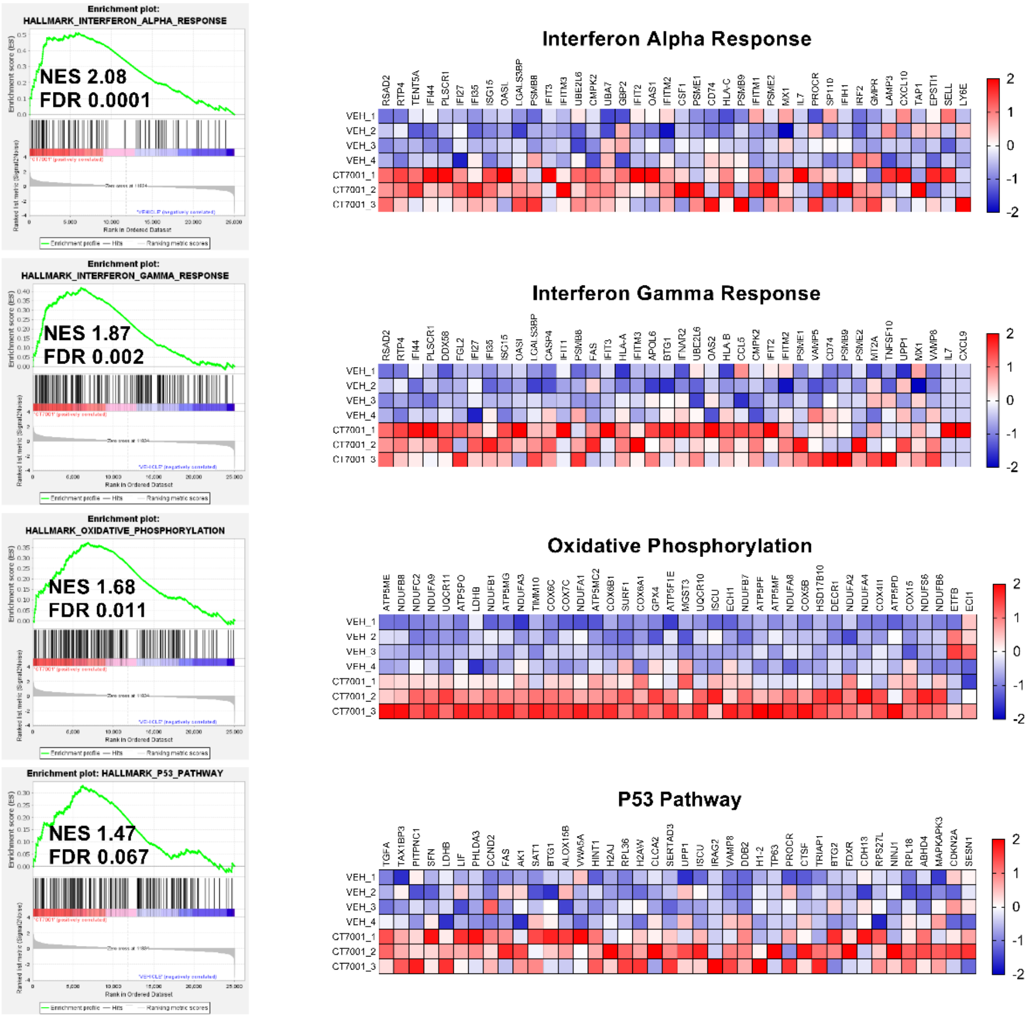


**Supplementary Figure S6. GSEA results for gene sets positively enriched in CT7001 treated tumours.** Enrichment plots and leading edge heatmaps (top 40 genes) are shown.
