## Supplemental Materials and Methods for "The CDK7 inhibitor CT7001 (Samuraciclib) targets proliferation pathways to inhibit advanced prostate cancer"

### SUPPLEMENTARY MATERIALS AND METHODS

#### Immunocytochemistry

LNCaP cells seeded on glass coverslips were allowed to adhere overnight before plating medium was replaced with starvation medium (phenol red-free RPMI-1640 supplemented with 5% charcoal-stripped FCS and 2mM L-glutamine). Cells were starved of androgens for 48 hours before treatment with DMSO or 10μM CT7001 in the absence or presence of androgen (10nM mibolerone) for 4 hours. Treated cells were washed with PBS, fixed with 4% paraformaldehyde in PBS for 15 minutes and permeabilized with 0.1% Triton-X diluted in PBS for 10 minutes at room temperature. Non-specific sites were blocked with 10% goat serum diluted in PBS for 1 hour at room temperature. Cells were stained with mouse anti-AR antibody (441) (sc-7305, Santa Cruz Biotechnology, 1:200) diluted in 1% BSA in PBS in a humidified chamber at 4°C overnight. The coverslips were washed 3 times in PBS for 5 minutes followed by staining with Alexa Fluor-594 secondary antibody (Invitrogen, 1:1000) diluted in 1% BSA in PBS for 1 hour at room temperature. Following three PBS washes, the coverslips were mounted on slides using DAPI-fluorescence mounting medium (Abcam). Fluorescence images were captured on a Nikon Eclipse E400 fluorescence upright microscope. For each treatment condition, 3 images were taken from 2 coverslips (6 images per condition). The experiment was repeated twice (n=2 biological repeats). Images were analysed using ImageJ v1.52a. A DAPI staining mask was used to define the nuclear regions of interest (ROI) for measuring mean nuclear AR fluorescence intensity.

#### Chromatin immunoprecipitation (ChIP)

LNCaP cells were grown in androgen depleted media for 72 hours before treatment for 4 hours with vehicle (DMSO) or 10 μM CT7001 in the presence of 10 nM mibolerone or equivalent vehicle (ethanol). Cells were fixed for 20 minutes with 1% formaldehyde, lysed, and sonicated to produce 300-1500 bp fragments. Chromatin immunoprecipitation was performed as described previously [1] using anti-AR (sc-7305, Santa Cruz Biotechnology) or rabbit mouse IgG at 10μg per sample and 100μL of Protein A Dynabeads (Invitrogen). DNA was isolated from samples using phenol:chloroform:isoamyl alcohol, washed, dried, and resuspended in nuclease-free water. Enrichment across three androgen-responsive elements (AREs) in the human PSA promoter, and PSA and FKBP5 enhancers was quantified by qPCR, with the primers listed in **Supplementary Table 2**. Data was calculated as percent input and normalized to non-specific control.

#### SUPPLEMENTARY TABLES

##### Supplementary Table 1. Primer sequences for RT-qPCR.

| Gene | Primer | Sequence (5'-3') |
|---|---|---|
| CDKN1A | Forward | CGATGGAACTTCGACTTTGTCA |
| CDKN1A | Reverse | GCACAAGGGTACAAGACAGTG |
| GADD45A | Forward | GAGAGCAGAAGACCGAAAGGA |
| GADD45A | Reverse | CACAACACCACGTTATCGGG |
| MDM2 | Forward | GAATCATCGGACTCAGGTACATC |
| MDM2 | Reverse | TCTGTCTCACTAATTGCTCTCCT |
| RRM2B | Forward | AGAGGCTCGCTGTTTCTATGG |
| RRM2B | Reverse | GCAAGGCCCAATCTGCTTTTT |
| BACTIN | Forward | GGCATCCTCACCCTGAAGTA |
| BACTIN | Reverse | GGTCATCTTCTCGCGGTTG |
| GAPDH | Forward | ATGGGGAAGGTGAAGGTCG |
| GAPDH | Reverse | GGGGTCATTGATGGCAACAATA |
| RPL19 | Forward | GCGGAAGGGTACAGCCAAT |
| RPL19 | Reverse | AGCAGCCGGCGCAAA |
| PSA | Forward | TTGTCTTCCTCACCCTGTCC |
| PSA | Reverse | AGCTGTGGCTGACCTGAAAT |

#### Supplementary Table 2. TaqMan low-density array qPCR assays

| Detector | Description | Response to androgens |
|---|---|---|
| 18S-Hs99999901_s1 | Eukaryotic 18S rRNA | Control |
| ABCC4-Hs00988717_m1 | ATP-binding cassette, sub-family C (CFTR/MRP), member 4 | Upregulated |
| ABHD2-Hs00199684_m1 | abhydrolase domain containing 2 | Upregulated |
| ACSL3-Hs00244853_m1 | acyl-CoA synthetase long-chain family member 3 | Upregulated |
| ALDH1A3-Hs00167476_m1 | aldehyde dehydrogenase 1 family, member A3 | Upregulated |
| ANKH-Hs00219798_m1 | ANKH inorganic pyrophosphate transport regulator | Upregulated |
| DBI-Hs01554584_m1 | diazepam binding inhibitor | Upregulated |
| DHCR24-Hs00207388_m1 | 24-dehydrocholesterol reductase | Upregulated |
| DNAJB9-Hs01052402_m1 | DnaJ (Hsp40) homolog, subfamily B, member 9 | Upregulated |
| ELL2-Hs01023022_m1 | elongation factor, RNA polymerase II, 2 | Upregulated |
| FKBP5-Hs01561006_m1 | FK506 binding protein 5 | Upregulated |
| GAPDH-Hs99999905_m1 | glyceraldehyde-3-phosphate dehydrogenase | Control |
| GNMT-Hs00219089_m1 | glycine N-methyltransferase | Upregulated |
| HMGCS1-Hs00940429_m1 | 3-hydroxy-3-methylglutaryl-CoA synthase 1 (soluble) | Upregulated |
| HPGD-Hs00960587_m1 | hydroxyprostaglandin dehydrogenase 15-(NAD) | Upregulated |
| IQGAP2-Hs00183606_m1 | IQ motif containing GTPase activating protein 2 | Upregulated |
| KLK2-Hs00428383_m1 | kallikrein-related peptidase 2 | Upregulated |
| KLK3-Hs02576345_m1 | kallikrein-related peptidase 3 | Upregulated |
| LIFR-Hs01123581_m1 | leukemia inhibitory factor receptor alpha | Upregulated |
| MAF-Hs00193519_m1 | v-maf avian musculoaponeurotic fibrosarcoma oncogene homolog | Upregulated |
| MTMR9-Hs00209995_m1 | myotubularin related protein 9 | Upregulated |

| | | |
|---|---|---|
| NDRG1-Hs00608387_m1 | N-myc downstream regulated 1 | Upregulated |
| NKX3-1-Hs00171834_m1 | NK3 homeobox 1 | Upregulated |
| ORM1-Hs01590791_m1 | orosomucoid 1 | Upregulated |
| PDIA5-Hs00895698_m1 | protein disulfide isomerase family A, member 5 | Upregulated |
| PGM3-Hs00985101_m1 | phosphoglucomutase 3 | Upregulated |
| PMEPA1-Hs00375306_m1 | prostate transmembrane protein, androgen induced 1 | Upregulated |
| PTPRM-Hs00267809_m1 | protein tyrosine phosphatase, receptor type, M | Upregulated |
| RAB27A-Hs00608302_m1 | RAB27A, member RAS oncogene family | Upregulated |
| RPLP0-Hs99999902_m1 | ribosomal protein, large, P0 | Control |
| SEPP1-Hs01032845_m1 | selenoprotein P, plasma, 1 | Upregulated |
| SLC35F2-Hs00213850_m1 | solute carrier family 35, member F2 | Upregulated |
| SORD-Hs00162091_m1 | sorbitol dehydrogenase | Upregulated |
| TBP-Hs99999910_m1 | TATA box binding protein | Control |
| TMPRSS2-Hs01120965_m1 | transmembrane protease, serine 2 | Upregulated |
| TPD52-Hs00893105_m1 | tumor protein D52 | Upregulated |

#### Supplementary Table 3. Primer sequences for ChIP-qPCR.

| Gene | Primer | Sequence (5'-3') |
|---|---|---|
| PSAprom | Forward | GTGCATCCAGGGTGATCTAGTAATT |
| PSAprom | Reverse | CACACCCAGAGCTGTGGAA |
| PSAenh | Forward | ACAGACCTACTCTGGAGGAAC |
| PSAenh | Reverse | AAGACAGCAACACCTTTTT |
| FKBP5enh | Forward | GCTCCCTCACACCAGATGACCA |
| FKBP5enh | Reverse | CAAATCCAACCCGAGACAGGTGTA |
